# The circular RNA landscape of human dorsal root ganglia and its association with opioid exposure

**DOI:** 10.1101/2025.02.26.640459

**Authors:** Maddy R. Koch, Jenna B. Demeter, Mark W. Shilling, Sascha R.A. Alles, Karin N. Westlund, Reza Ehsanian, June Bryan I. de la Peña

**Affiliations:** Department of Anesthesiology and Critical Care Medicine, University of New Mexico Health Sciences Center, Albuquerque, New Mexico, USA

**Keywords:** circRNA, opioids, DRG, nociceptors, pain signaling, transcriptomics

## Abstract

Opioids are among the most widely prescribed treatments for pain; however, prolonged use leads to adverse effects, including reduced analgesic efficacy (tolerance) and paradoxically heightened pain sensitivity (opioid-induced hyperalgesia, OIH). Neurons that detect noxious stimuli within the dorsal root ganglion (DRG), referred to as nociceptors, mediate both the beneficial and maladaptive effects of opioids. Although post-transcriptional regulation is critical for DRG function, the role of circular RNAs (circRNAs)—an evolutionarily conserved and highly stable class of RNA—in nociceptive processes remains largely unexplored in humans. Further, how opioids might alter the circRNA landscape of human DRG (hDRG) is unknown. To address this gap, we performed high-coverage RNA sequencing on hDRG tissue obtained from opioid-positive organ donors and compared these profiles with those from age- and sex-matched opioid-negative controls. The circRNA expression profiles were analyzed using the CIRI2/CIRIquant pipeline, and parallel measurements were made for the linear transcriptome (e.g. mRNA). Our data revealed a significant overall decrease in circRNA abundance in the opioid-exposed group. Among the top differentially expressed circRNAs (FDR ≤ 0.05) were *circSH3D19*, *circSMARCA5*, *circHLA-A*, and *circAMY2B*, with an additional 39 circRNAs (p ≤ 0.005) altered in opioid-exposed tissue. To explore potential interactions with the linear transcriptome, we constructed a competing endogenous RNA (ceRNA) network using established pipelines and databases (circAtlas, miRanda, TargetScan, PITA, and miRDB). Gene Ontology enrichment analysis of predicted mRNA targets of these circRNAs identified overrepresented pathways related to neuronal development, synaptic signaling, inflammatory processes, and pain perception. These findings suggest that circRNAs may play a key regulatory role in the DRG’s response to opioid exposure and modulation of pain. Future studies will investigate the spatial and temporal dynamics and functional and behavioral effects of these circRNA.

## Introduction

Chronic pain is a major global health burden, affecting millions of individuals and contributing significantly to disability and healthcare costs.^1–4^ In the United States alone, the economic burden of pain is estimated to range from $560 to $635 billion annually, surpassing the combined costs of cancer and diabetes.^5^ Despite its prevalence, effective pain management remains a persistent challenge, emphasizing the urgent need for deeper investigations into molecular pain mechanisms and the development of novel therapeutic strategies.

Opioids are potent analgesics widely used to manage moderate to severe pain; however, their prolonged use is associated with potential for abuse, diminished analgesic efficacy (tolerance) and a paradoxical increase in pain sensitivity, known as opioid-induced hyperalgesia (OIH).^6–9^ These maladaptive effects complicate pain management, leading to dose escalation, dependence, and contributing to the ongoing opioid abuse epidemic. The dorsal root ganglion (DRG), a key hub for pain signal transmission, houses neurons that detect noxious stimuli (nociceptors) that play a critical role in both the beneficial (analgesic) and detrimental (tolerance, OIH) effects of opioids.^10–12^ Opioid receptors are abundantly expressed in human and rodent DRG, implicating these neurons in opioid-induced adaptations.^13^ Notably, in rodents, morphine tolerance and OIH are dependent on MOR signaling in DRG nociceptors.^14^ Given their unique anatomical structure, with long axonal projections extending to peripheral tissues and the spinal cord, DRG neurons rely heavily on post-transcriptional RNA regulatory mechanisms.^15–18^ While opioid-induced molecular adaptations in DRG neurons have been investigated at the transcriptional and epigenetic levels,^19,20^ the role of post-transcriptional regulatory mechanisms, particularly circular RNAs (circRNAs), remain largely unexplored.

circRNAs are a class of non-coding RNAs generated through back-splicing, forming a covalently closed loop structure that renders them resistant to exonuclease degradation.^21,22^ These molecules are highly stable and exert numerous regulatory functions, including serving as microRNA (miRNA) sponges to limit their activity, interacting with RNA-binding proteins, and direct binding to the 3’ untranslated region (UTR) of their parent genes.^23,24^ circRNAs are enriched in the nervous system, where they play essential roles in neurodevelopment, synaptic plasticity, and neurodegenerative diseases.^25^ Recent work has revealed diverse and multifaceted ways in which circRNAs regulate gene expression in neuronal tissue beyond simple miRNA and RBP sponging. For instance, in the rodent brain, *circDlc1(2)* physically interacts with both miR-130b-5p and various mRNAs associated with glutamatergic signaling to synergistically repress expression of these genes at the synapse.^26^ In *Drosophila*, neuronal splicing factor Muscleblind (MLB) can promote circularization of its own locus to form a negative feedback loop *in-cis*.^27^ Further, a short-hairpin RNA (shRNA) targeted at the backsplice junction of *circMbl* induces developmental lethality and impairments, implicating unique functions of *circMbl in-trans*. These studies highlight the diverse roles of circRNAs and critical functions they play in the nervous system.

In rodents, circRNAs undergo alterations in the sensory neurons of the DRG in response to various pain conditions, including peripheral neuropathy and arthritis.^28–30^ For example, in a rodent model of neuropathic pain (chronic constriction injury) significantly modified expression of 374 circRNAs in the DRG was observed, with the top 50 interacting with pain-related microRNAs.^30^ Emerging evidence also shows that circRNAs are dynamically regulated in response to opioid exposure. For instance, studies in mice have shown that *circOPRM1*, a circRNA derived from the mu-opioid receptor (*OPRM1*) gene, is dysregulated in the brain and spinal cord following repeated opioid treatment.^31^ However, opioid-induced circRNA dysregulation in the DRG, particularly in humans, and its role in pain processing and neuronal adaptations remain largely unexplored, highlighting a critical gap in our understanding of opioid-induced molecular changes in nociceptors.

In this study, we investigated the impact of opioids on the circRNA–mRNA landscape in human dorsal root ganglion (hDRG) tissue obtained from ethically consented organ donors with verified opioid exposure, as indicated by a positive opioid result in their terminal toxicology report. Age- and sex-matched opioid-negative donors served as controls. Using high-coverage RNA sequencing and an established circRNA analysis pipeline (CIRI2/CIRIquant), we identified significant alterations in circRNA expression in opioid-exposed hDRG tissue. Additionally, we constructed a competitive endogenous RNA (ceRNA) regulatory network and performed Gene Ontology (GO) enrichment analysis to identify pathways associated with pain processing and opioid adaptation. Our results provide the first comprehensive landscape of circRNAs in hDRG and are the first to demonstrate that opioid exposure induces significant alterations in circRNA expression in human nociceptors. These findings suggest that circRNAs may play a critical regulatory role in mediating opioid-related adaptations within the DRG. Elucidating these molecular changes may reveal novel therapeutic targets for mitigating opioid-induced adverse effects and enhancing pain management strategies.

## Materials and Methods

### Human dorsal root ganglion (hDRG) tissue acquisition

DRG tissue was obtained from ethically consented organ donors at the University of New Mexico Hospital in collaboration with New Mexico Donor Services. This study was conducted in compliance with Institutional Review Board (IRB) guidelines and was approved by the University of New Mexico Health Sciences Center Human Research Review Committee (Approval numbers #21-412 and #23-205). Research procedures adhered to the ethical principles outlined in the Declaration of Helsinki. Donor demographic details are summarized in **Table 1**.

**Table 1:**
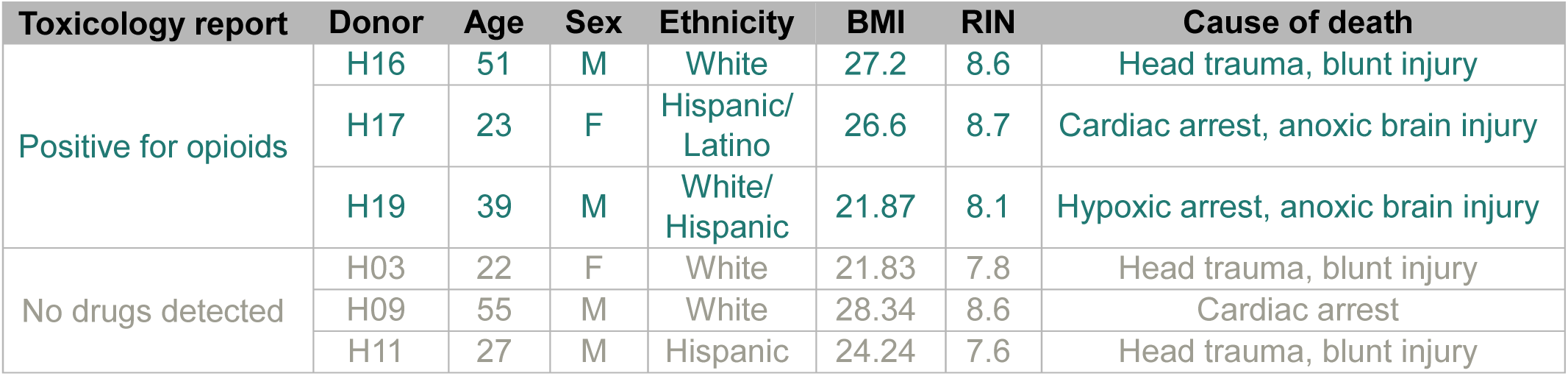
Donor information for the six organ donors whose DRG tissue was used for transcriptomic profiling. The table includes toxicology results, age, sex, ethnicity, BMI, RNA integrity number (RIN), and cause of death.

### RNA sequencing

Total RNA was isolated from frozen hDRG tissue (n = 3 per condition: opioid-exposed and control) using the Monarch Total RNA Miniprep Kit (New England Biolabs) following the manufacturer’s protocol. The quantity and purity of isolated RNA were initially assessed using a NanoDrop spectrophotometer (Thermo Fisher Scientific), ensuring OD 260/280 ≥ 2.0 and OD 260/230 ≥ 1.8, indicating no or minimal contamination. The integrity of the RNA was further evaluated using the 2100 Bioanalyzer (Agilent), with samples achieving an RNA Integrity Number (RIN) ≥ 7.5 deemed suitable for library construction.

Total RNA with rRNA removed underwent library construction. The libraries were sequenced on a NovaSeq X Plus Sequencing System (Illumina) using a 150-bp paired-end read strategy. This sequencing approach provided comprehensive coverage, generating > 14 Gb of raw data per sample, which resulted in > 45 million raw read pairs for each sample. This depth of coverage ensured robust quantification of long transcripts and accurate detection of circRNA and linear RNA (mRNA and lncRNA). Phred scores—Q20 > 97% and Q30 > 92%—for all samples indicated that sequencing was of high quality and that data was suitable for further analysis.

### Bioinformatic analysis

#### Quality control

Raw data underwent filtering of paired reads when either read 1) was contaminated with an adapter, 2) had > 10% uncertain nucleotides (N) (rarely), or 3) was of low quality (Q < 5 for > 50% of nucleotides) (very rarely). Poly-G tails—which occur in two-color chemistry systems when a black G is repeatedly called after synthesis has terminated—were trimmed using BBDuk when ≥ 20 bp. Quality was ensured by several metrics using FastQC.

#### Alignment, quantification, and differential expression

CIRIquant^32^ was used as a comprehensive tool for circRNA junction site prediction, alignment and quantification of back-splice junction (BSJ) and forward-splice junction (FSJ) reads to a circRNA index, edgeR-based differential expression of circRNAs, and basic annotation of circRNAs. The GRCh38, release 112 Ensembl reference genome was used for all analyses. circRNAs were predicted using the default program, CIRI2. Age- and sex-matched pairs of an opioid-exposed and a control donor were utilized for the analysis as follows: 1) H17 and H03, 2) H16 and H09, and 3) H19 and H11. circRNA with p ≤ 0.005 and |log2FC| ≥ 1 were considered differentially expressed circRNAs (DECs).

For alignment and quantification of linear transcripts, HISAT2 and stringTie were employed. The edgeR generalized linear model quasi-likelihood pipeline was used for differential expression analysis of transcripts, with opioid exposure (positive or negative) and age- and sex-matched pair as factors in the design matrix. Transcripts with p ≤ 0.01 and |log2FC| ≥ 1 were considered differentially expressed transcripts (DETs). This Ensembl-HISAT2-StringTie-edgeR pipeline was used for consistency with CIRIquant pipeline and given the fact it has been ranked as the overall most accurate pipeline for identifying DEGs in a recent RNA-seq benchmarking study.^33^

#### Determination of detected and expressed circRNA and comparison by tissue

To determine the number of circRNAs detected in various tissues, we referenced by tissue type the number of entries in the circAtlas database, which had inclusion criteria of ≥ 2 independent BSJ reads.^34^ To determine the number of “detected” circRNA in our hDRG comparably, we counted only circRNA that had ≥ 1 count in ≥ 2 samples. Meanwhile, circRNAs with ≥ 2 counts in ≥ half (3/6) of our hDRG tissue samples were considered “expressed” and utilized for further analyses. CPM was averaged per group (opioid-exposed vs. control) for each circRNA when relevant.

#### Calculation of circRNA CPM

We calculated the counts per million (CPM) of circRNAs using the linear transcript library size (total number of counts) rather than the circRNA library size. This allowed us to contextualize the abundance of circRNA in a manner comparable to the entire transcriptome (which is predominantly linear) and determine whether differences in circRNA abundance are associated with opioid exposure.

#### PCA and heatmaps

Principal component analysis (PCA) of hDRG tissue samples was conducted in base R using raw counts that underwent trimmed mean of M-values (TMM) normalization in edgeR. Scaling was used for the linear transcript PCA to account for the wide inter-transcript variation in counts/ expression levels. The heatmaps of circRNA or linear transcript expression in opioid-exposed vs.

control hDRG tissue samples were generated using pheatmap, with log2CPM (for circRNAs) or log2TPM (for DETs) as input and scaling by circRNA/ transcript to generate Z-scores for each sample.

#### Over-representation analysis

Gene Ontology (GO) Biological Process (BP) over-representation analysis (ORA) was conducted using clusterProfiler, using an “interesting” gene set coupled to a “background” gene set (all genes that passed filterbyExpr in edgeR) as input. For the ORA of the parent genes of circRNAs at least moderately expressed in hDRG (average CPM ≥ 1 among the 6 samples), the interesting gene set was the parent genes of these expressed circRNA, identified as all genes that fall within the BSJ genomic coordinates per CIRIquant. For the ORA of the parent genes of the DECs, the interesting gene set was the parent genes of the DECs, identified as the genes that span the BSJ genomic coordinates. For the DET ORA, the interesting gene set was the genes corresponding to the DETs.

A threshold of p ≤ 0.05 was used to define significantly enriched terms. While significantly enriched terms representing a variety of biological processes were identified, key representative terms were selected along the themes of neurogenesis and morphogenesis (including neuron development and differentiation as well as neuron projection) and signaling (synaptic; membrane potential and ion flux; and transmembrane, sensory, pain, and stress) given the ubiquity of these themes in the significantly enriched term sets as well as their relevance to neuronal activity and pain modulation.

#### Competitive endogenous RNA (ceRNA) network construction

For each DEC in the opioid-exposed vs. control group, miRNAs predicted to have ≥ 1 binding site by (v3.3a),^35^ TargetScan (v7.0),^36^ or the probability of interaction by target accessibility (PITA; v2.1.2)^37^ tool where accessed through circAtlas.^38^ This compiled list was then filtered using the miRNA expression profile of hDRG samples available in the NOCICEPTRA database.^39^ Only miRNAs with ≥100 reads per million (RPM) in this dataset were included. Using data from miRBD,^40^ predicted mRNA targets of these miRNAs (target score ≥ 80) were compiled. For the ceRNA network, these targets were filtered by mRNA DETs in the opioid-exposed vs. control group. For the ceRNA-associated ORA of circRNA-specific, miRNA-mediated mRNA targets, the interesting gene set was predicted mRNA targets with ≥1 CPM in ≥ half (3/6) samples in our hDRG tissue dataset. Conversion between RefSeq gene IDs, Ensembl gene IDs, and Ensembl transcript IDs was performed using biomaRt.^41^ The network was visualized using CytoScape (v3.10.3).^42^

## Results

### circRNAs are abundantly expressed in human DRG tissue and are enriched in genes related to neuronal development, synaptic function, and pain

Although rodent studies are available, to our knowledge, there is no published data characterizing the circRNA landscape of hDRG. To address this gap, we performed high-coverage RNA sequencing of hDRG tissue to capture total long transcripts and mapped, annotated, and quantified the expression of circRNAs using the CIRI2/CIRIquant pipeline. We identified 20,034 unique circRNA species expressed in ≥ two independent samples (**Fig. 1A**). Using tissue-specific publicly available data compiled in circAtlas, we found that this number is comparable to other tissues, most notably the spinal cord (22,272 unique circRNAs). The most abundant circRNAs in hDRG include *CDR1as* (all), *circHIPK3* (exon 2), and *circN4BPL2* (exons 3-6) (**Fig. 1B**). An over-representation analysis (ORA) of parent genes revealed that circRNAs in hDRG are enriched in genes related to neuronal development, differentiation, projection, synaptic function, ion flux, sensation, pain, and stress (**Fig. 1C**). The vast majority of these circRNAs are exonic (94.5%), although intronic (4.82%) are also abundant (**Fig. 1D**). The most common biotype of circRNA parent transcripts was protein-coding (94.34%) followed by lncRNA (8.1%), with significant overlap between these categories due to the presence of circRNAs with multiple parent transcripts (**Fig. 1E**). Almost all circRNAs with a pseudogene (4.82%), miRNA (1.04%), or other small non-coding RNA (1.64%) parent transcripts also had a protein-coding parent. Of exonic parent genes, most only produced a single circRNA (54.9%), although 7.52% produced five or more (**Fig. 1F**). The median putative length of exonic circRNAs was 2,677 nucleotides (nts), with the greatest frequency between 500-1500 nts (**Fig. 1G**). Collectively, these findings provide the first comprehensive landscape of circRNAs in the hDRG and suggest that circRNAs may perform regulatory functions similar to those in other neuronal tissues such as the brain and spinal cord.

**Figure 1.**
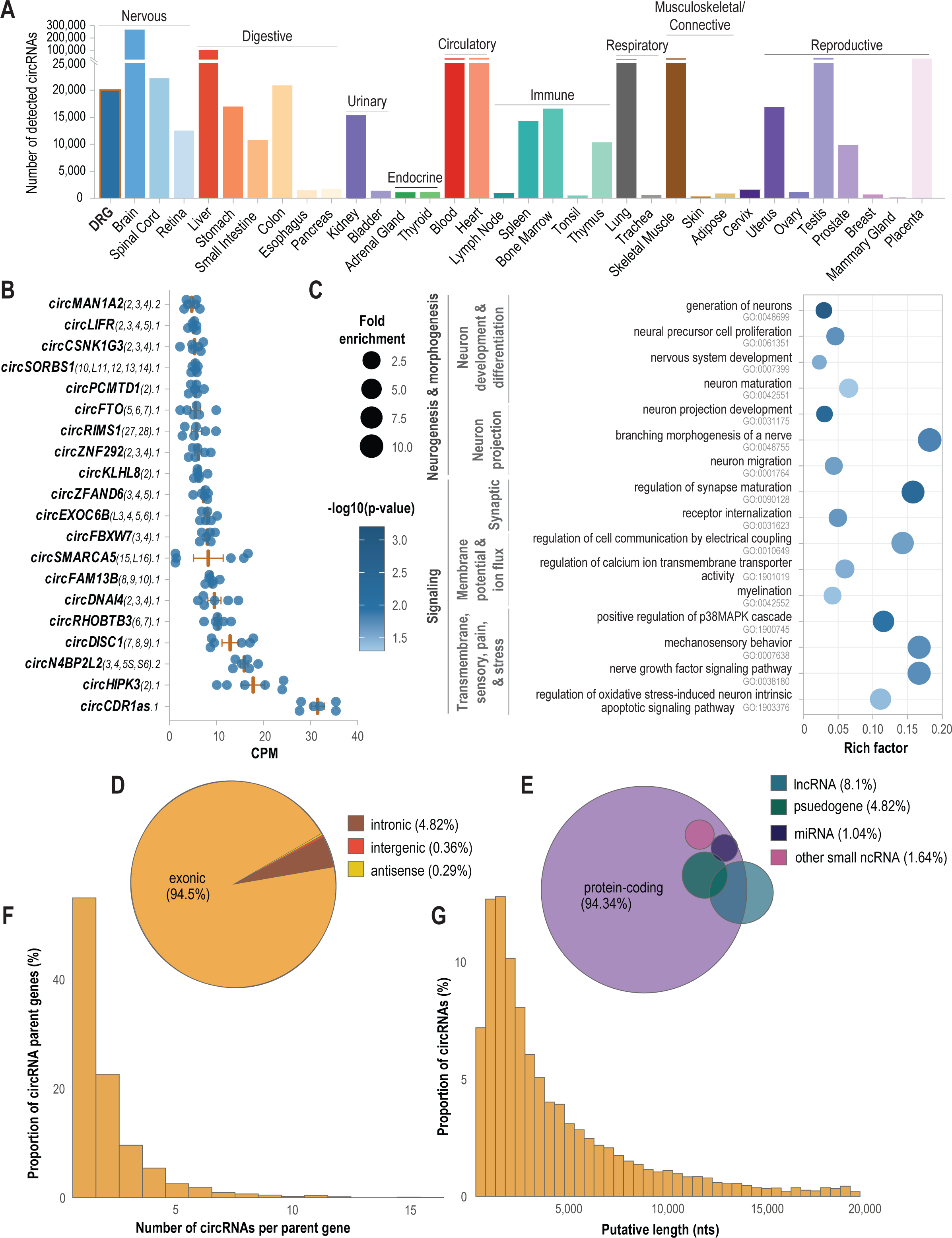
Characterization of circRNA landscape of the human DRG. **(A)** Comparison of the number of circRNAs with ≥ 1 count in ≥ 2 independent samples in our dataset compared to the number of circRNAs other tissues in circAtlas. (**B**) Expression (CPM) of circRNAs with the highest abundance, sorted from bottom to top. (**C**) Select GO terms enriched in circRNA parent genes, divided into categories relevant to sensory neurons and nociceptive processes. (**D**) Proportion (%) of exonic, intronic, and intergenic circRNAs. (**E**) Euler plot of circRNA parent gene biotype. Other small non-coding RNA includes scaRNA, snoRNA, and snRNA. (**F**) Histogram of the number of circRNAs per parent gene. (**G**) Histogram of exonic putative circRNA length, assuming circRNAs contain all exons (spliced) within the circRNA genomic region.

### Opioid-exposure is associated with alterations in the circRNA landscape of hDRG tissue

To investigate the impact of opioids on circRNA expression in hDRG tissue, samples from three donors (1 female, 2 male) that tested positive for opioids during a terminal toxicology report (hereafter referred to as “opioid-exposed”) were compared to age- and sex-matched controls. Further information on donors can be found in **Table 1**. First, using the average expression of circRNAs per group, we found a global decrease in density of circRNA abundance in opioid-exposed tissue (Kolmogorov-Smirnov test, p < 0.05) (**Fig. 2A**). Notably, only a limited number of circRNAs were unique to each group (60 in controls and 30 in opioid-exposed samples; **Fig. 2B**). A three-dimensional principal component analysis (PCA) of circRNA expression showed distinct clustering of opioid-exposed donors (**Fig. 2C**). Differential expression analysis identified 25 upregulated and 18 downregulated circRNAs in the opioid-exposed group (|log₂FC| ≥ 1;p ≤ 0.005). The top five DECs were *circSH3D19* (exons 15–17; log₂FC = –6.69, p = 1.74E–09), *circSMARCA5* (exons 15–16; log₂FC = –3.56, p = 1.93E–07), *circHLA-A* (exons 1–8; log₂FC = 9.07, p = 9.45E–07), *circAMY2B* (exons 1–6; log₂FC = –7.30, p = 3.23E–06), and *circNTRK2* (exons 3–6; log₂FC = –3.67, p = 6.29E–05) (**Fig. 2D**). A heatmap of circRNA expression showed that upregulated and downregulated circRNAs largely clustered into distinct groups (**Fig. 2E**). Finally, over-representation analysis of the parent genes of these DECs revealed significant enrichment of GO terms related to neurogenesis, morphogenesis, synaptic function, and ion influx (**Fig. 2F**). Together, these findings indicate that opioid exposure is associated with both global and specific alterations in the circRNA landscape of the hDRG, indicating a potential role for circRNAs in opioid-induced neuronal adaptations.

**Figure 2.**
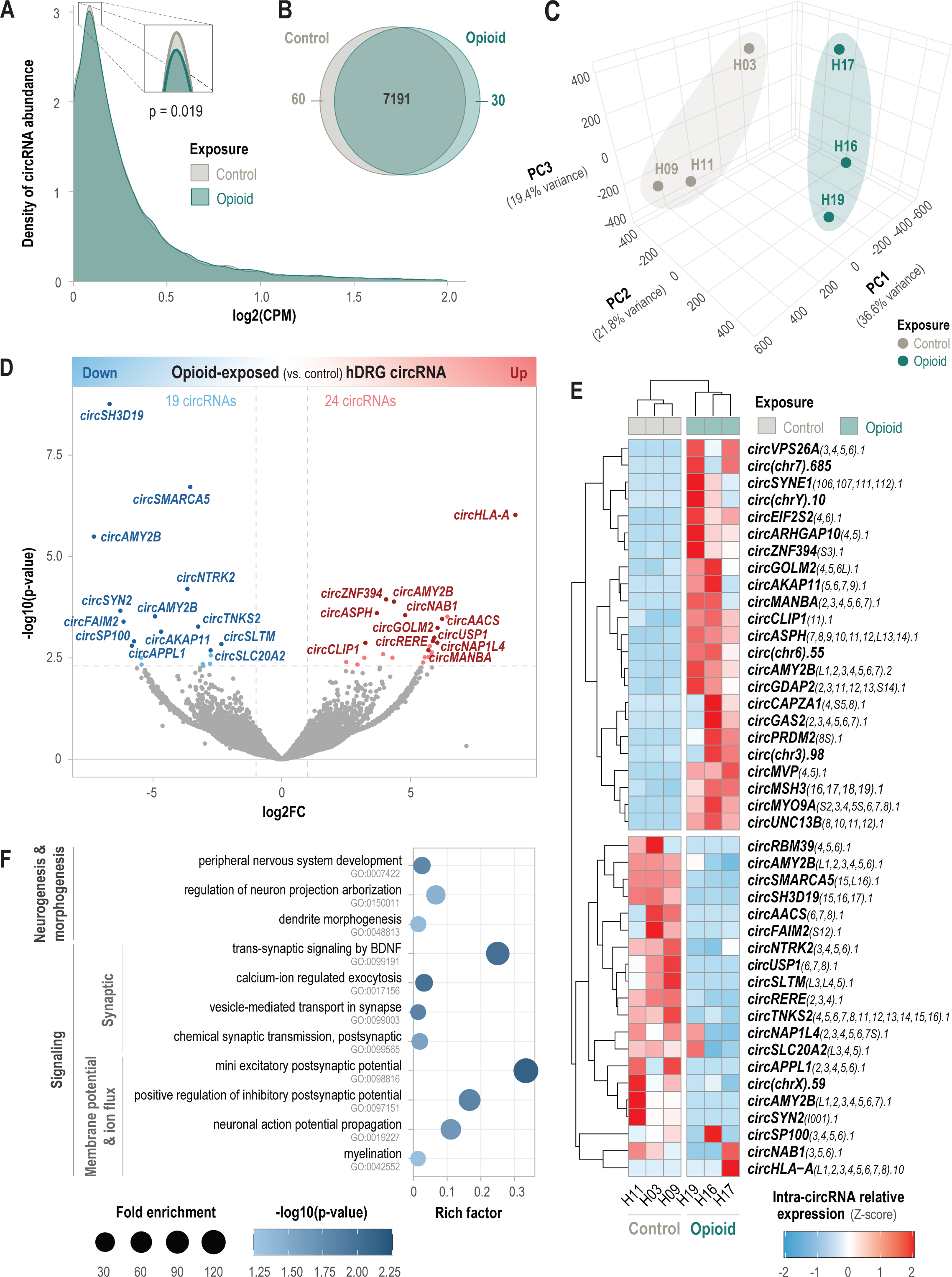
Circular transcriptome profile of hDRG associated with opioid exposure. **(A)** Density plot of average circRNA abundance for opioid-exposed (blue) and control (grey) donors, filtered by circRNAs with ≥ 2 counts in ≥ half the samples. (**B**) Venn diagram showing the number of circRNAs with ≥ 2 counts in ≥ half the samples unique to each and common among opioid-exposed and control groups. (**C**) Three-dimensional PCA plot representing the circular transcriptome profile of hDRG from opioid-exposed and control donors. (**D**) Volcano plot demonstrating up- and downregulated circRNA in hDRG from opioid-exposed donors relative to control donors. Upregulated transcripts (log_2_FC ≥ 1, p ≤ 0.005) are indicated in red while downregulated transcripts (log_2_FC ≤ -1, p ≤ 0.005) are indicated in blue. (**E**) Heatmap of the cross-sample relative expression of the top differentially expressed transcripts (DETs). Samples and genes are clustered hierarchically, with respective dendrograms to the side. (**F**) Select GO terms significantly (p ≤ 0.05) enriched in DETs, divided into categories relevant to sensory neurons and nociceptive processes. Size indicates fold enrichment and color gradient indicates -log10(p-value).

### Opioid-exposure is associated with alterations in the long linear transcriptome of hDRG tissue

To investigate linear transcriptomic changes associated with opioid exposure in hDRG tissue, we quantified and compared hDRG tissue long transcript expression (mRNA and lncRNA) between opioid-exposed donors and age and sex-matched controls using a HISAT2-stringTie-edgeR pipeline. Although a PCA of transcript expression did not show clustering of opioid-exposed donors (**Fig. 3A**), 176 transcripts were downregulated and 173 upregulated in the opioid-exposed group (|log₂FC| ≥ 1; p ≤ 0.01) (**Fig. 3B**). The top five DETs included F-box and leucine rich repeat protein 5 (*FBXL5*)-209 (log₂FC = 4.83, p = 7.38E-05), adenosine deaminase RNA specific (*ADAR*)-222 (log₂FC = -7.36, p = 1.03E-04), tenascin XB (*TNXB*)-208 (log₂FC = 2.21, p = 2.97E-04), secreted frizzled related protein 5 (*SFRP5*)-201 (log₂FC =-1.66, p=3.17E-04), and nexilin F-actin binding protein (*NEXN*)-202 (log₂FC = -6.94, p = 3.67E-04). Interestingly, of the top 50 DETs, 6 of them encoded different isoforms of *HLA-DQ* (A1-206, A1-205, B1-201, B1-AS1-201, B1-202, and B1-205), a member of the human leukocyte antigen (HLA) class II complex, all significantly upregulated (**Fig. 3C**). Other immune-related genes (e.g. *IL12RB1*, *ADAR*, *NFKB1*) are also represented among these transcripts. Similar to circRNA parent genes, an ORA of these transcript genes revealed enrichment in GO biological process terms related to neurogenesis and morphogenesis, synaptic function, ion influx, and inflammation (**Fig. 3D**). Taken together, these findings indicate opioid exposure is associated with alterations in transcripts related to neuronal function and inflammation in hDRG tissue.

**Figure 3.**
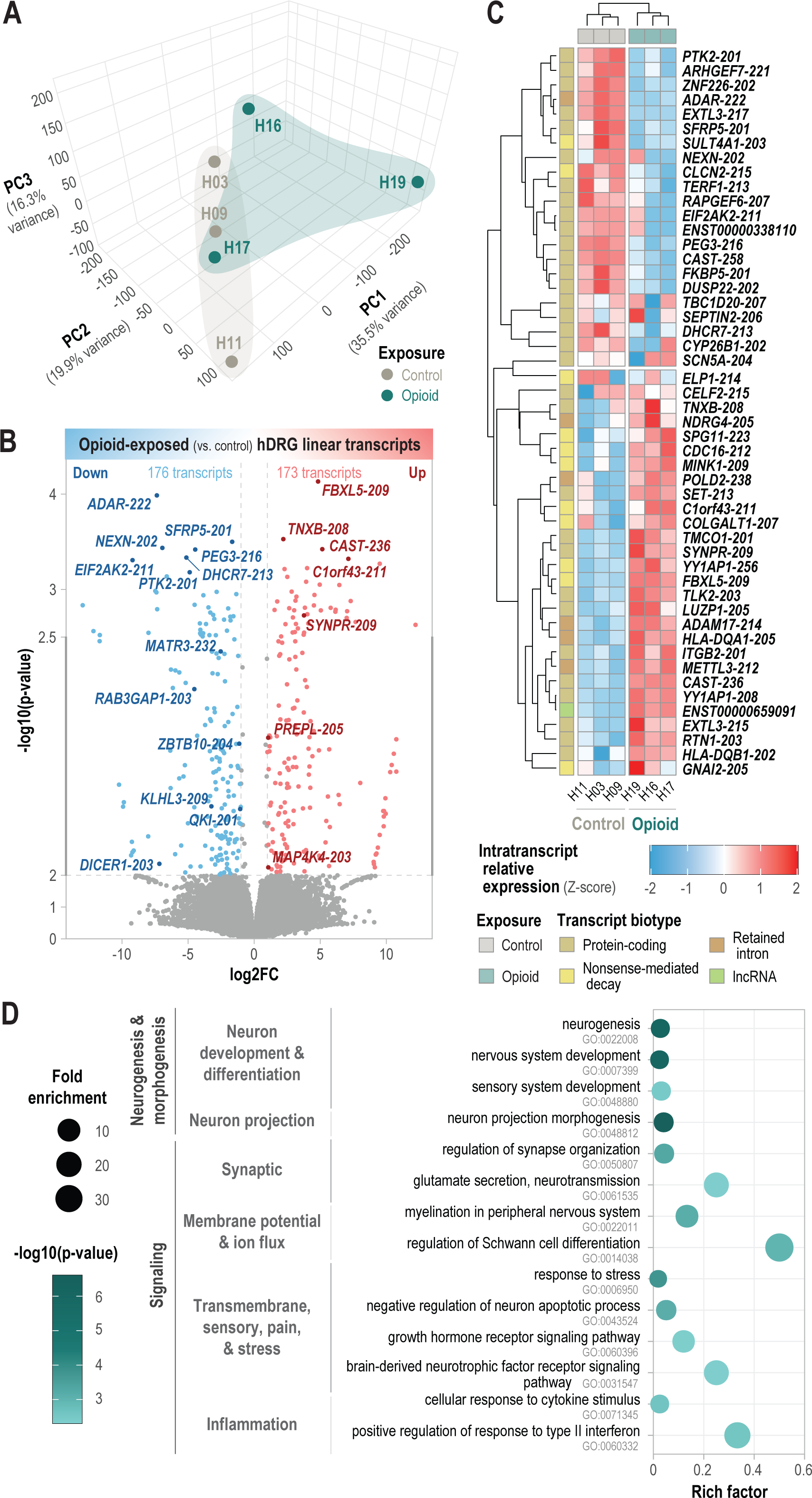
Linear transcriptome profile of hDRG associated with opioid exposure. **(A)** Three-dimensional PCA plot representing the linear transcriptome profile of hDRG from opioid-exposed and control donors. (**B**) Volcano plot demonstrating up- and downregulated linear transcripts in hDRG from opioid-exposed donors relative to control donors. Upregulated transcripts (log_2_FC ≥ 1, p ≤ 0.01) are indicated in red while downregulated transcripts (log_2_FC ≤ - 1, p ≤ 0.01) are indicated in blue. The top 10 DETs, as well as DETs with ≥ 3 circRNA inputs in the ceRNA network are labeled. (**C**) Heatmap of the cross-sample relative expression of the top 50 differentially expressed transcripts (DETs). Samples and genes are clustered hierarchically, with respective dendrograms to the side. Type of exposure (opioid vs. control) and transcript biotype (protein-coding, nonsense-mediated decay, retained intron, or lncRNA) are indicated by color for samples and transcripts, respectively. (**D**) Select GO terms significantly (p ≤ 0.05) enriched in DETs, divided into categories relevant to sensory neurons and nociceptive processes. Size indicates fold enrichment and color gradient indicates -log10(p-value).

### Opioid-exposure associated circRNAs may form a ceRNA network to alter expression of genes related to pain in hDRG tissue

Sponging of miRNAs to regulate expression of their target mRNA transcripts is a well characterized function of circRNAs.^21^ To investigate potential miRNA-mediated regulatory relationships among circRNAs (**Fig. 2D**) and linear transcripts (**Fig. 3B**) differentially expressed in opioid-exposed hDRG tissue, we constructed a ceRNA network using miRNA binding prediction data from miRanda, targetscan, PITA (circRNAs) and miRBD (mRNAs) (**Fig. 4A**). Because our high-coverage RNA sequencing pipeline did not capture small non-coding RNAs, we filtered the predictions by miRNAs experimentally shown to be expressed in hDRG neurons in the NOCICEPTRA database.^39^ The resulting ceRNA network contains 18 circRNAs, 28 miRNAs, and 44 mRNAs, with 119 edges (**Fig. 4B**). The circRNAs with the most targets in this network are *circSLC20A2* (5 miRNAs targeting 13 genes), *circZNF394* (6 miRNAs targeting 10 genes), and *circGAS2* (4 miRNAs targeting 10 genes). Conversely, the mRNAs receiving the most inputs in the network are *KLHL3* (2 miRNAs targeted by 6 circRNAs), *MAP4K4* (1 miRNA targeted by 6 circRNAs), and *ZBTB10* (4 miRNA targeted by 4 circRNAs). A heatmap of intragene expression shows upregulated and downregulated network DETs largely cluster separately (**Fig. 4C**). Next, we supposed that although a given DEC may cause small effects on individual transcript expression, this might amount to a significant alteration across a wider functional network. To investigate this, we performed an ORA on predicted miRNA-mediated mRNA targets of individual DECs, regardless of whether these transcripts were significantly altered in our dataset (**Fig. 4D**). GO terms significantly (p ≤ 0.05) enriched in miRNA-mediated mRNA targets of DECs contained substantial overlap, with neuron development enriched across all circRNAs analyzed. Overall, miRNA-mediated mRNA targets were enriched in GO terms related to neuronal development and differentiation, projection, synaptic function, and sensation, pain, and stress. Taken together, these findings suggest potential miRNA-mediated mechanisms through which opioid-associated circRNAs might contribute to dysregulation of specific transcripts and broader functional networks in hDRG tissue, which may contribute to nociceptor plasticity.

**Figure 4.**
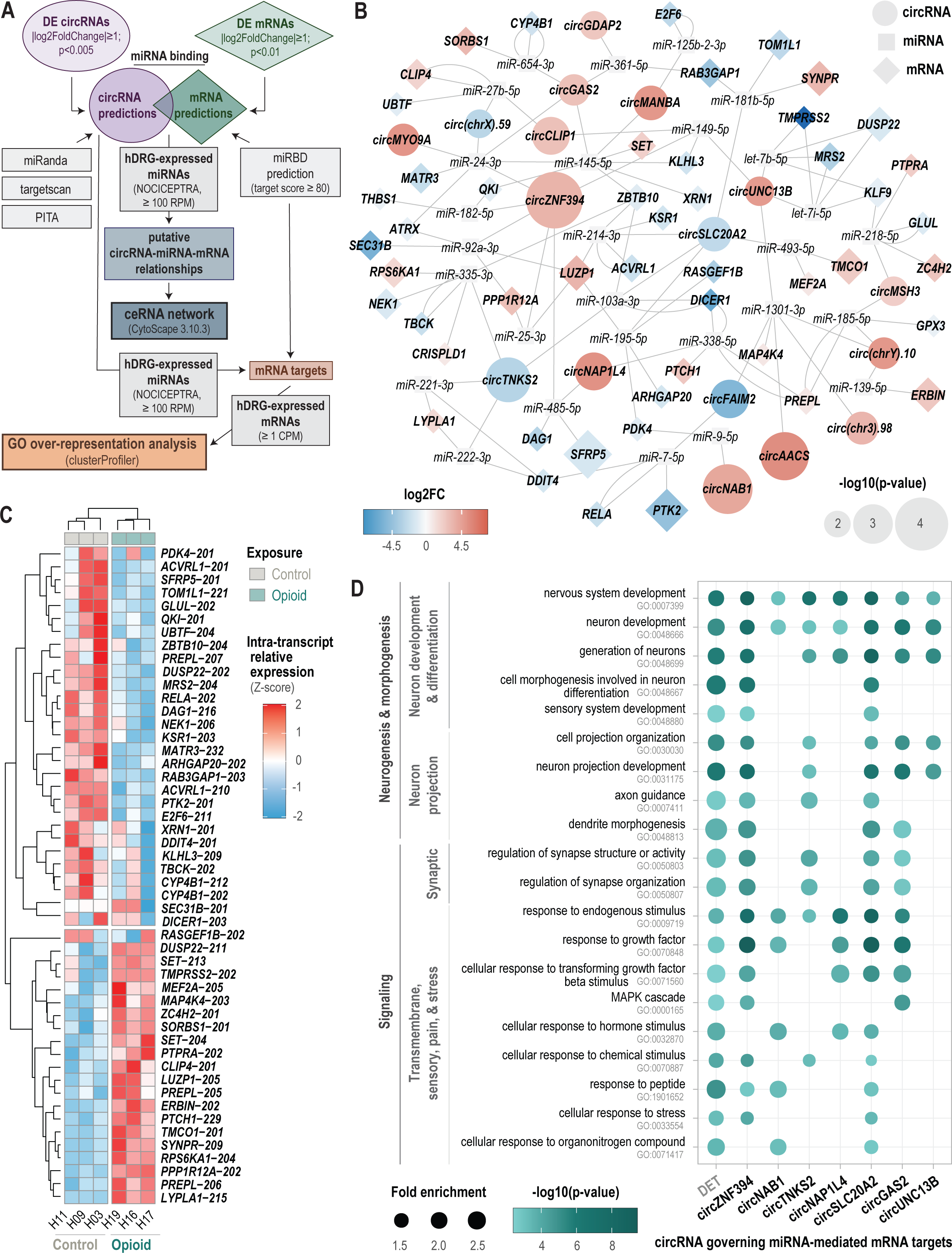
Competitive endogenous RNA (ceRNA) network of differentially expressed circRNAs (DECs) and mRNA transcripts in opioid-exposed hDRG tissue. **(A)** Graphical summary of ceRNA network construction and ORA of hDRG-expressed miRNA targets of select DECs. (**B**) ceRNA network of DECs and differentially expressed transcripts (DETs). Shape indicates RNA type (circles = circRNA, squares = miRNA, and diamonds = mRNA), size indicates -log10(p-value), and color indicates log_2_FC, with red upregulated and blue downregulated. Edges indicate predicted binding between RNAs. Multiple edges between a given miRNA/mRNA pair indicate multiple DETs for that gene contain binding sites for the miRNA. (**C**) Heatmap of the cross-sample relative expression of network DETs. Samples and genes are clustered hierarchically, with respective dendrograms to the side. (**D**) Select GO terms significantly (p ≤ 0.05) enriched in total DRG-expressed miRNA targets of select DECs. Size indicates fold enrichment and color gradient indicates -log10(p-value).

## Discussion

In this study, we present the first comprehensive landscape of circRNAs in human DRG tissue and demonstrate that opioid exposure is associated with a global decrease in circRNA abundance along with specific alterations in circRNA expression. In parallel, opioid use also induced significant transcriptomic changes, particularly affecting genes involved in immune and neuronal signaling. Our network analysis further suggests that a circRNA-miRNA-mRNA post-transcriptional regulatory mechanism may underlie these transcriptomic alterations.

CircRNAs are well-established as key post-transcriptional regulators in the central nervous system, where they are enriched at synapses and modulate neuronal activity. Although circRNAs have been investigated in rodent DRG, studies in human DRG are lacking. Significant species differences are reported between rodent and human DRG transcriptomes.^43–44^ Thus, our data provide a critical, clinically relevant perspective on the circRNA landscape in human DRG and its potential role in somatosensation and pain signaling.

Our analysis revealed that the overall abundance of circRNAs in the hDRG is comparable to that observed in the spinal cord (Fig. 1A). The most abundant circRNAs, including well-characterized species such as *CDR1as*, *circHIPK3*, and *circSMARCA5*, have previously been implicated in synaptic plasticity and neuroprotection. For example, *CDR1as* is a highly conserved circRNA containing 74 miR-7 binding sites, known to modulate glutamatergic signaling and synaptic plasticity in the rodent hippocampus and cortical neurons.^45–47^ Similarly, *circHIPK3* has been reported to protect against inflammation and neuronal apoptosis following spinal cord injury,^48^ while *circSMARCA5* is thought to function as a competing endogenous RNA (ceRNA) across multiple cancer types.^49^ Notably, circRNAs in hDRG tissue are enriched in GO terms related to neuronal development, synaptic function, membrane potential regulation, and sensory/pain pathways, underscoring their potential regulatory roles in peripheral nociception.

Opioids are potent analgesics; however, their adverse effects, particularly tolerance and OIH, represent a major challenge. Sensory neurons in the DRG, which abundantly express opioid receptors, contribute significantly to these phenomena.^13,14,50^ Given that opioids modulate circRNAs in the brain and spinal cord, we hypothesized that similar mechanisms might occur in the hDRG. Indeed, our data shows that opioid-exposed donors exhibited a marked decrease in circRNA abundance compared to controls. This could potentially be due to the fact that opioids decrease neuronal activity and circRNAs are produced in response to increases in neuronal activity.^51^ Interestingly, while PCA based on circRNA expression clearly distinguished opioid-exposed donors (Fig 2C), PCA of the overall linear transcriptome did not show such separation (Fig. 3A), suggesting that opioid exposure preferentially affects the circRNA transcriptome. Differential expression analysis identified 19 downregulated and 24 upregulated circRNAs. The top DEC was *circSH3D19*, derived from the SH3 domain containing 19 linear transcript. Although the function of its parent gene has not been characterized in the peripheral nervous system, it is known to interact with members of the ADAM (a disintegrin and metalloproteinase) family, which are involved in peripheral nerve regeneration and nociception.^52–54^ Interestingly, *circSMARCA5*, mentioned above as one of the most highly expressed circRNAs, was the second most downregulated in opioid-exposed tissue. However, the function of the majority of DECs have not been characterized, presenting opportunities for further research. The parent genes of opioid-responsive circRNAs are enriched for functions related to neuronal development and signaling, indicating a potential role for circRNAs in modulating DRG circuitry and nociceptive transmission.

In addition to circRNA alterations, our analysis of the linear transcriptome revealed significant changes in immune and neuronal signaling genes. The top upregulated transcript was *FBXL5*, which is critical to iron hemostasis,^55^ a process which is dysregulated in neurons by opioid exposure.^56,57^ The top downregulated was *ADAR*, which is downregulated by repeated morphine exposure in rodent spinal neurons, thereby triggering an inflammatory response through increased secretion of double stranded RNA, ultimately leading to hyperalgesia.^58^ Also among the top DETs are *TNXB*, which has been associated with mechanical allodynia and nociceptor hypersensitivity in mice,^59^ *CAST* (calpastatin), a regulator of axonal degradation following peripheral nerve injury,^60^ and *DHCR7* (7-dehydrocholesterol reductase), which has been shown to regulate sensory neuron survival via its effect on cholesterol metabolism in Schwann cells.^61^ Enrichment analysis of these transcripts revealed overrepresentation of pathways related to peripheral nervous system development, synaptic function, ion influx, and inflammatory responses, and ion influx, which are all processes involved in peripheral nociception.^62^ Interestingly, these terms are also consistent with the roles of the parent genes of opioid-responsive circRNAs.

To explore the potential regulatory role of circRNAs in mediating these transcriptomic changes, we constructed a ceRNA network integrating DECs, miRNA binding predictions, and DETs. We utilized hDRG miRNA expression data from NOCICEPTRA database. Our network highlights one of the top 10 DECs, *circZNF394*, which is predicted to compete with one the top 10 DETs, *SFRP5*, for binding of *miR-485-5p*. The circRNAs with the most miRNA targets, *circSLC20A2*, *circZNF394*, and *circGAS2*, are all uncharacterized but are prime candidates for further investigation. Although the mRNA with the most inputs, *KLHL3*, a ubiquitin ligase, does not have studies directly implicating it in nociception, its well-established target WNK1,^63,64^ is known to play a role in peripheral neuropathy through regulation of NKCC1 and KCC2 in sensory neurons.^65,66^ Another mRNA receiving many circRNA inputs in the network is *MAP4K4*, which has been shown to regulate inflammation and the stress response of the DRG.^67,68^ Although the network analysis is limited by the absence of single-cell and matched miRNA data, it provides a valuable framework for understanding how circRNAs may contribute to opioid-induced dysregulation of gene expression in the hDRG.

Despite these and other limitations (e.g. small sample size, variability in human donor tissues, and lack of comprehensive history of opioid use) our findings offer critical insights into the molecular adaptations associated with opioid exposure in the hDRG. Future studies will focus on functional validation of circRNA roles in primary hDRG neuronal cultures and sensory neuron from *in vivo* chronic pain models, as well as longitudinal assessments of circRNA dynamics with repeated opioid exposure. Integration of proteomic and epigenetic data would also be helpful to construct a comprehensive multi-omic model of opioid-induced adaptations. Additionally, targeted interventions based on circRNA modulations through antisense oligonucleotide (ASO) may pave the way for novel, non-opioid analgesic strategies.

In summary, our study highlights the importance of circRNAs as potential regulators of gene expression related to pain processing in the DRG following opioid exposure. By using human tissue, we enhance the clinical relevance of our findings and provide novel insights into circRNA functions human primary sensory neurons. This database and the networks described herein lay the foundation for future mechanistic studies aimed at elucidating the direct effects of opioids on circRNA expression and nociceptive signaling in the hDRG.

